# ABL1 Kinase plays an important role in spontaneous and chemotherapy-induced genomic instability in multiple myeloma

**DOI:** 10.1101/2022.05.12.491727

**Authors:** Subodh Kumar, Srikanth Talluri, Jiangning Zhao, Chengcheng Liao, Lakshmi B. Potluri, Leutz Buon, Shidai Mu, Jialan Shi, Chandraditya Chakraborty, Yu-Tzu Tai, Mehmet K. Samur, Masood A. Shammas, Nikhil C. Munshi

## Abstract

Genomic instability contributes to cancer progression and is at least partly due to dysregulated homologous recombination. Here, we show that an elevated level of ABL1 kinase overactivates the HR pathway and causes genomic instability in multiple myeloma (MM) cells. Inhibiting ABL1 with either shRNA or a pharmacological inhibitor (nilotinib) inhibits HR activity, reduces genomic instability, and slows MM cell growth. Moreover, inhibiting ABL1 rescues the HR dysregulation and genomic instability caused by melphalan, a chemotherapeutic agent used in MM treatment, and increases melphalan’s efficacy and cytotoxicity *in vivo* in a subcutaneous tumor model. In these tumors, nilotinib inhibits endogenous as well as melphalan-induced HR activity. These data demonstrate that inhibiting ABL1 using the clinically approved drug nilotinib reduces MM cell growth, promotes genome stability, increases the cytotoxicity of melphalan (and similar chemotherapeutic agents), and can potentially prevent or delay progression in MM patients.

## Introduction

Genomic instability or adaptability is a common feature of cancer cells^1–6^ that enables them to acquire mechanisms that confer greater growth and resistance to treatment^7–11^. In multiple myeloma, a cancer of the plasma cells, tumor cells continue to acquire genomic changes over time, and this genomic instability is linked to disease progression^6,11–13^. A fundamental problem is that most chemotherapeutic agents are DNA damaging agents. Although such treatment can kill a large fraction of cancer cells, it is also likely to increase chromosomal instability in surviving cancer cells, possibly resulting in the emergence of drug-resistant clones and subsequent relapse. In fact, having an increased rate of DNA repair allows a cancer cell to survive treatment and appears to be associated with chemotherapy resistance in several malignancies^14–17^. Particularly in relapsed MM patients, melphalan resistance is associated with increased rates of DNA repair^18^.

Homologous recombination (HR), which accurately repairs DNA by copying the missing information from the homologous sequence, maintains genomic integrity and stability^19–22^. Dysregulation or hyperactivation of HR can lead to aberrant forms of HR such as non-allelic homologous recombination, homeologous recombination, and microhomology-based recombination. These aberrant forms are associated with a variety of genomic aberrations^23–26^. For example, microhomology-mediated break induced replication can cause chromothripsis, which is characterized by massive genomic rearrangements in a localized area of the genome^35^. Chromosomal fragments and whole chromosomes produced by defective DNA repair and genomic instability can get encapsulated in a lipid bilayer to form micronuclei^32^, which can induce further DNA damage through aberrant DNA repair within the micronucleus^33–34^ In fact, spontaneously elevated HR in multiple myeloma contributes to the acquisition of genomic changes over time as well as the emergence of drug resistance^11^. Thus, dysregulated HR in a cancer cell not only repairs DNA breaks but can also produce new breaks (to initiate unscheduled or unnecessary repair), mediate genomic rearrangements (through its aberrant forms), and increase overall genomic instability and chemotherapy resistance.

ABL1 tyrosine kinase is associated with various aspects of DNA damage repair^36^. The protein c-ABL phosphorylates RAD51, which is the central component of HR repair^37^. Likewise, BCR-ABL1 kinase-mediated phosphorylation of RAD51 promotes aberrant HR with a possible role in the progression of chronic myeloid leukemia (CML)^38^. In fact, tyrosine kinase inhibitors (such as imatinib and nilotinb) targeting ABL1 are currently used in the treatment of CML patients, with nilotinib being the drug of choice for chronic myeloid leukemia in the chronic phase^38^. Although the BCL-ABL1 fusion gene has not been reported in multiple myeloma, a recent study suggests that ABL1 signaling plays a role in plasma cell survival^40^. We therefore evaluated the role of ABL1 kinase in endogenous and chemotherapy-induced genomic instability and growth of MM cells. We show that elevated expression of ABL1 contributes to overactivation of HR and genomic instability in MM, and its inhibitor nilotinib inhibits chemotherapy-induced genomic damage and instability, while increasing cytotoxicity.

## MATERIALS AND METHODS

### Cell lines and patient samples

Normal human fibroblasts, non-cancerous human bone marrow stromal cell line (HS-5) and human multiple myeloma cell lines MM.1S, and RPMI8226 were purchased from American Type Tissue Culture Collection (Rockville MD). KMS-12-BM, AMO1, U266, NCI-H929 and JJN3 and were purchased from DSMZ (Braunschweig, Germany). All cell lines used were confirmed to be free of mycoplasma and cultured in RPMI1640 growth medium, which contained 10% fetal bovine serum (HyClone, South Logan, UT) as reported previously^11,41–43^.

Blood samples from healthy donors were collected, and the peripheral blood mononuclear cells (PBMCs) isolated by Ficoll-Paque gradient (GE Healthcare, Boston, MA, USA). Informed consent was obtained in accordance with the Helsinki Declaration and the Review Board of the Dana-Farber Cancer Institute.

### Reagents

Nilotinib and melphalan (Sigma Aldrich, Saint Louis, MO) were dissolved in DMSO and diluted in cell culture medium to the desired concentrations prior to use. Anti-RAD51 antibody, catalogue # SC-8349, was purchased from Santa Cruz, Dallas, TX. Antibodies against GAPDH (catalogue # mAb2118), HP1 (catalogue # 261) and γH2A.X (Ser139, catalogue #2577) were from Cell Signaling Technology, Inc., Danvers, MA.

### Western Blotting

Total cell lysates were prepared using the Thermo Fisher Scientific Cell Lysis Buffer (Thermo Fisher Scientific) supplemented with a protease and phosphatase inhibitor cocktail (Thermo Fisher Scientific). Equal amounts of proteins were resolved by SDS-polyacrylamide gel electrophoresis and transferred to a nitrocellulose membrane. Immunoblotting was performed using the following antibodies: β-actin (Santa Cruz), γ-H2AX and RAD51 (Cell Signaling).

### Lentiviral particles and infections

Suppression of ABL1 was achieved using lentivirus-based shRNAs (Sigma Aldrich). Briefly, cells were transduced with lentiviral particles in the presence of 8-μg/mL polybrene and selected in 2-μg/mL puromycin. Before the experiments, dead cells were eliminated using a Ficoll-Pague gradient (GE Healthcare, Boston, MA, USA) and live cells were allowed to recover for 24 h.

### Immunofluorescence staining and microscopy

For immunofluorescence staining, 20,000 cells were suspended in 100 μl of 1.5% BSA in PBS, cytospun onto glass slides at 1000 rpm for 5 min, fixed in 4% paraformaldehyde for 10 min, washed with PBS and permeabilized with 0.1% Triton X-100 containing 1.5% BSA in PBS for 10 min. Cells were then incubated overnight at 4°C with primary antibodies against γ-H2AX (mouse monoclonal anti-γH2AX antibody) and RAD51 (rabbit polyclonal anti-RAD51 antibody, Santa Cruz). Cells were washed and incubated with the appropriate fluorescein-conjugated secondary antibodies (Alexa Fluor 594-labeled goat anti-mouse IgG from Abcam for γ-H2AX; Alexa Fluor 488-labeled goat anti-rabbit IgG from Abcam for RAD51). After washing, cells were mounted under coverslips with Prolong Antifade reagent containing DAPI (Invitrogen). Images were acquired with Yokogawa Spinning Disk Confocal/TIRF System with a 63x oil objective lens.

### Cell Viability Assays

Cells were seeded at 2000–5000 per well in 96-well plates, treated as indicated for 72 h and assessed for cell viability using the Cell Titer-Glo® Luminescent Cell Viability Assay (Promega) or Cell Counting Kit-8 (CCK-8) assay (Sigma Aldrich) according to the manufacturer’s protocol. Viability was expressed as percentage (± SEM) of three or more independent biological replicates relative to control-treated cells.

### Recombination Assays

Homologous recombination (HR) activity was assessed using either an *in vitro* HR assay described previously^42^ or a homologous strand exchange assay as reported^42^.

### Detection of DNA breaks

The extent of DNA breaks was assessed by evaluating MM cells for the levels of γ-H2AX. Expression of γ-H2AX was measured by Western blotting using anti-γ-H2AX (Ser139, antibody # 2577; Cell Signaling Technology, Inc., Danvers, MA) as reported by us previously^42,44,47^.

### Evaluating impact on genomic instability

To evaluate the impact on genomic instability, the cells were cultured in the presence or absence of nilotinib, melphalan or a combination of both, and were evaluated for micronuclei, which is a marker of an unstable genome^30–32^, as reported by us previously^42–47^. Genomic instability was also assessed using single nucleotide polymorphism (SNP) arrays (Affymetrix). Briefly, cells were cultured in the presence or absence of nilotinib. An aliquot of cells was harvested and frozen at the beginning of the experiment (Day 0) to be used as a reference or baseline genome. The acquisition of new copy number alterations in culture relative to the reference cells was monitored using SNP arrays as described by us previously^47^.

### Impact of nilotinib/melphalan in a subcutaneous tumor model

Five-week-old male CB-17 SCID mice were purchased from Charles River Laboratories (Wilmington, MA) and maintained following the guidelines of the Institutional Animal Care and Use Committee (IACUC). All experimental procedures were approved by the IACUC and the Occupational Health and Safety Department of the Dana-Farber Cancer Institute, Boston, MA, USA. MM.1S cells (3.0×10^6^ in 100 μl saline) were injected subcutaneously into the interscapular area of each mouse. Following the appearance of tumors (~7-10 days), the mice were treated with normal saline containing a working dilution of solvent DMSO, nilotinib (50 mg/kg; 5 days per week) and/or melphalan (2 mg/kg; twice per week), injecting intraperitoneally. Tumor sizes were measured regularly, and animals sacrificed when tumors reached 2 cm^3^ in volume or when paralysis or major compromise in their quality of life occurred. Tumors were removed and evaluated for HR activity.

## Results

### ABL1 contributes to elevated homologous recombination and more DNA breaks in MM cells

Compared to normal plasma cells, *ABL1* was upregulated across all stages of plasma cell dyscrasias (p = 0.004, dataset - GSE2568) (**Figure 1A**). Concordantly, ABL1 protein expression was also upregulated in MM cell lines compared to undetectable levels in normal PBMCs (**Figure 1B)**. Since ABL1 regulates RAD51 recombinase through phosphorylation, we investigated the HR activity in MM cell lines. ABL1 was inhibited using nilotinib or shRNA-mediated knock-down and the impact on HR evaluated using a functional HR activity assay. We observed significant inhibition of HR activity in both MM cell lines with shRNA and in a dose-dependent manner with nilotinib (**Figure 1C, D**). Moreover, nilotinib also inhibited RAD51 phosphorylation and its ability to form foci in MM cells (**Figure 1E, F**). The nilotinib treatment also reduced the number of DNA breaks, which are the first stage of the HR process, as evidenced from reduced γH2AX expression and fewer γH2AX foci. Consistently, ABL1 knockdown also reduced the number of spontaneous DNA breaks in MM cells (**Figure 1G**).

**Figure 1.**
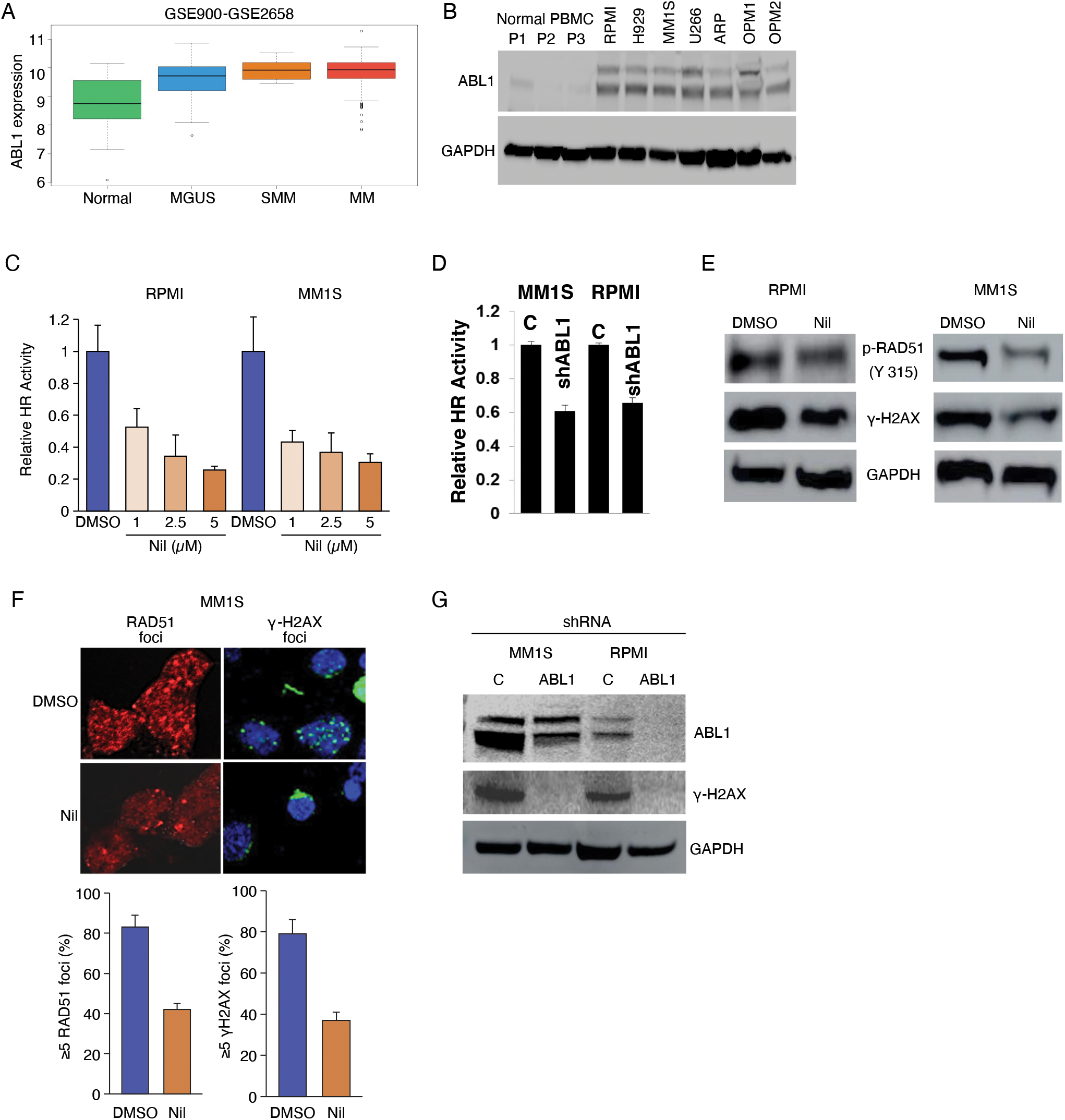
ABL1 kinase is overexpressed and contributes to dysregulation of HR activity and genomic integrity in MM. **(A)** Relative expression (Log2) of *ABL1* in the GSE2658 dataset (N=16, MGUS=22, SMM=24 and MM=73) between normal and MM samples. **(B)** Western blot showing ABL1 expression in normal PBMC samples and MM cell lines. **(C-D)** HR activity in MM cell lines treated with DMSO (D; control), the ABL1 inhibitor nilotinib (Nil; 5 μM) for 48 hrs (panel C) or shRNAs (C, control; sh-ABL1, ABL1-shRNA) (panel D). **(E)** MM cell lines treated with DMSO or nilotinib (Nil; 5 μM) for 48 h were evaluated for phosphorylation of RAD51 (Y315) and DNA breaks (by investigating levels of Ͱ-H2AX) using Western blotting and **(F)** microscopy. **(G)** ABL1 was suppressed using shRNAs (C, control; sh-ABL1, ABL1-shRNA) and the cells were evaluated for DNA breaks (by investigating levels of Ͱ-H2AX) using Western blotting.

### Inhibition of ABL1 reduces acquisition of new genomic changes over time in MM

Given that inhibiting ABL1 reduced HR activity, we next investigated whether it could reduce genomic instability. We inhibited ABL1 in MM cells by gene knockdown or nilotinib and investigated the impact of these modulations on the number of micronuclei, a marker of ongoing genomic instability and rearrangements^30–32^. ABL1 inhibition in both MM cell lines significantly reduced (~50%; P<0.05) the number of micronuclei in three independent experiments, indicating reduced genomic instability (**Figure 2A, B**). To further investigate genomic instability on a finer scale, we cultured MM cells in the presence or absence of nilotinib for three weeks and investigated the acquisition of new copy number events relative to “day 0” cells (serving as the baseline genome), using single nucleotide polymorphism arrays (Affymetrix). Images of copy number events acquired on chromosomes 4 and 9 are shown as example (Figure 2C, left panel). Over the whole genome, the acquisition of copy number change events in nilotinib-treated cells was reduced to ~55% of control cells (**Figure 2C**). These data confirm that ABL1 inhibition reduces genomic instability in MM cells and suggest that it could reduce the rate of genomic evolution in MM patients.

**Figure 2.**
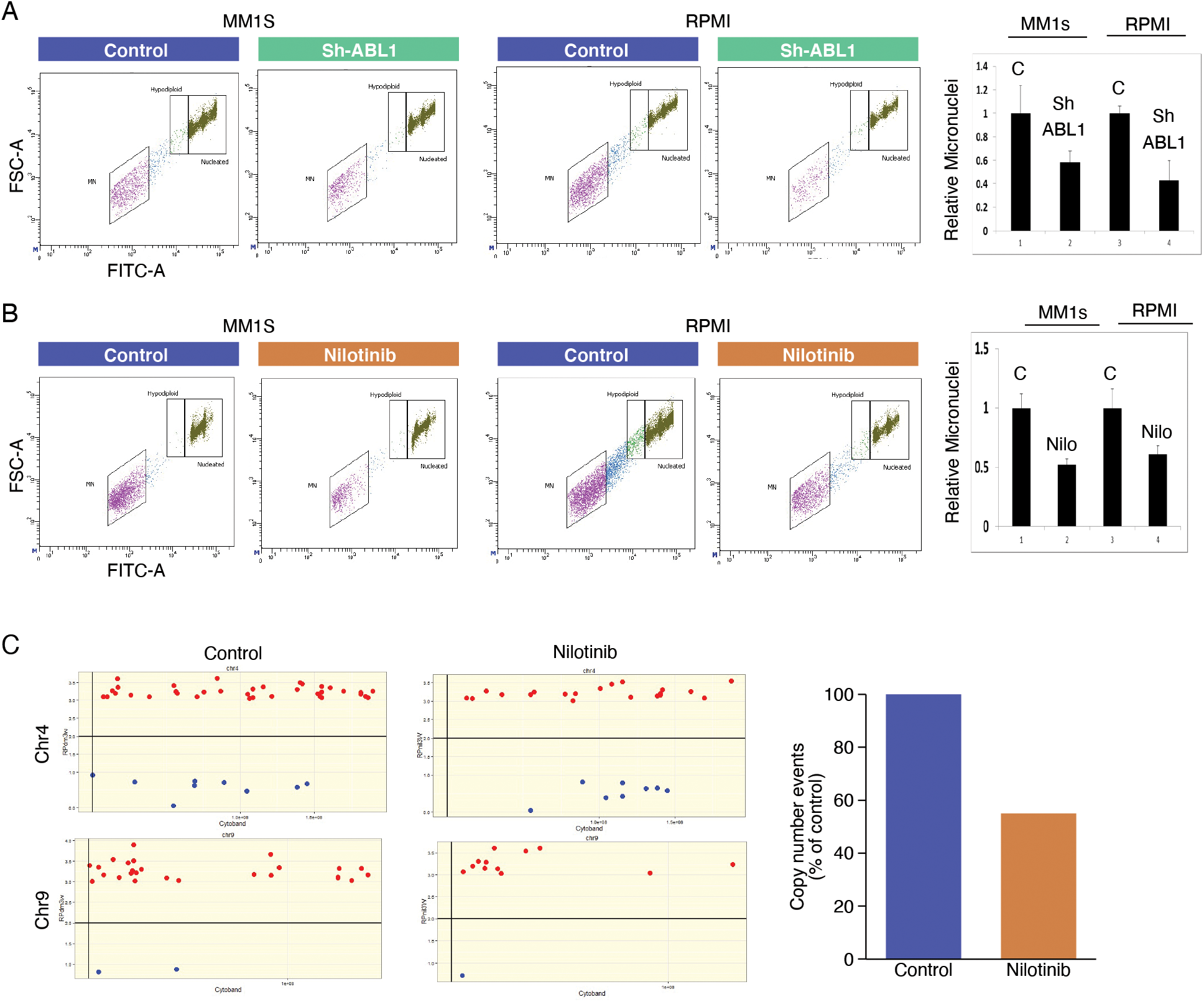
Inhibition of ABL1 by shRNA or nilotinib reduces genomic instability in MM cells. **(A-B)** ABL1 was inhibited in MM cell lines (MM1S and RPMI8226) either using shRNA (C, control; sh-ABL1, ABL1-shRNA) or by culturing in the presence of DMSO or 5 μM ABL1 inhibitor (nilotinib) for 48 hrs, and cells were evaluated for genomic instability using the micronucleus assay. Flow cytometry images of micronuclei and bar graphs showing percent micronuclei (with error bars representing SDs of three independent experiments) for each cell line are shown. **(C)** RPMI8226 cells were treated with DMSO only (control) or nilotinib for 21 days, and acquisition of new copy number events relative to the baseline genome of “day 0” cells was evaluated using SNP6.0 arrays (Affymetrix). Representative image of copy number events (amplifications, red dots; deletions, blue dots) on chromosomes 4 and 9 and bar graph showing copy events over whole genome are shown.

### Inhibition of ABL1 reduces melphalan-induced HR activity in MM cells

Melphalan is a chemotherapeutic agent that causes extensive DNA damage. In fact, treatment with melphalan increases RAD51 expression and HR activity in MM cells in a dosedependent manner (**Figure 3A, B**), likely due to the cells attempting to repair their genome. To investigate if ABL1 inhibition can reverse melphalan-induced HR activity, thus increasing melphalan cytotoxicity, we treated MM cells with nilotinib (N) and melphalan (M), alone as well as in combination, and evaluated the impact on HR activity. Whereas melphalan significantly increased HR activity, nilotinib not only inhibited endogenous HR activity but also reversed melphalan-induced HR activity in MM cells (**Figure 3C**).

**Figure 3.**
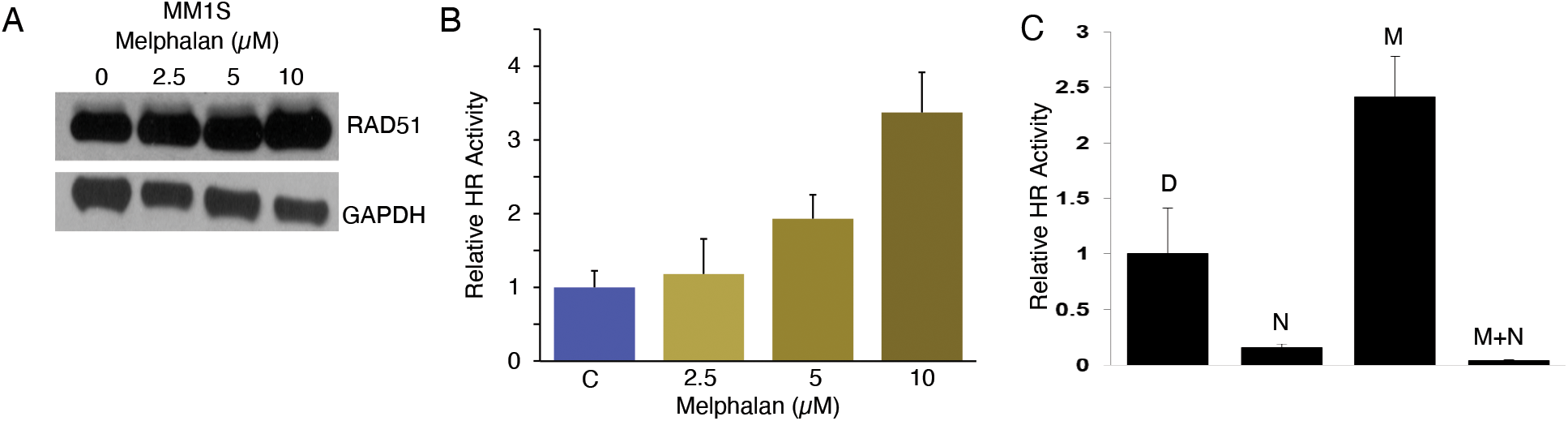
Melphalan-induced HR activity in MM cells is reversed by nilotinib. **(A)** MM1S cells were treated with different concentrations of melphalan and evaluated for RAD51 expression by Western blotting and **(B)** HR activity. **(C)** HR activity assessed in MM1S cells treated with nilotinib (2.5 μM), melphalan (5 μM) and a combination of both drugs for 48 hrs.

### Inhibition of ABL1 reduces melphalan-induced DNA breaks and genomic instability in MM cells

Given that ABL1 inhibition reduces melphalan-induced HR activity, we next investigated whether melphalan also induces genomic instability and whether that could be reversed by ABL1 inhibition. Briefly, we treated MM cells with nilotinib (2.5 μM), melphalan (5 μM) and a combination of both drugs for 48 hrs and evaluated the live cell fractions for DNA breaks (by measuring γ-H2AX level) and genomic instability (using the micronucleus assay). Melphalan was associated with a massive increase in the number of DNA breaks in the live MM cell fraction, and nilotinib almost completely reversed this (**Figure 4A**). Concurrently, melphalan increased the number of micronuclei by ~1.8 – 2-fold in all three cell lines, whereas nilotinib significantly reduced the number of both the basal and melphalan-induced micronuclei in all MM cell lines tested (**Figure 4B, C**). We used the same copy number assay described above on MM1S cells cultured in the presence or absence of nilotinib and melphalan for three weeks and found that, over the whole genome, melphalan led to a >12-fold increase in the number of copy number events whereas nilotinib not only inhibited the spontaneous but also the melphalan-induced acquisition of copy number events in MM cells (**Figure 4D**). Altogether, these data show that ABL1 inhibition reduces spontaneous as well as melphalan-induced genomic instability in MM cells.

**Figure 4.**
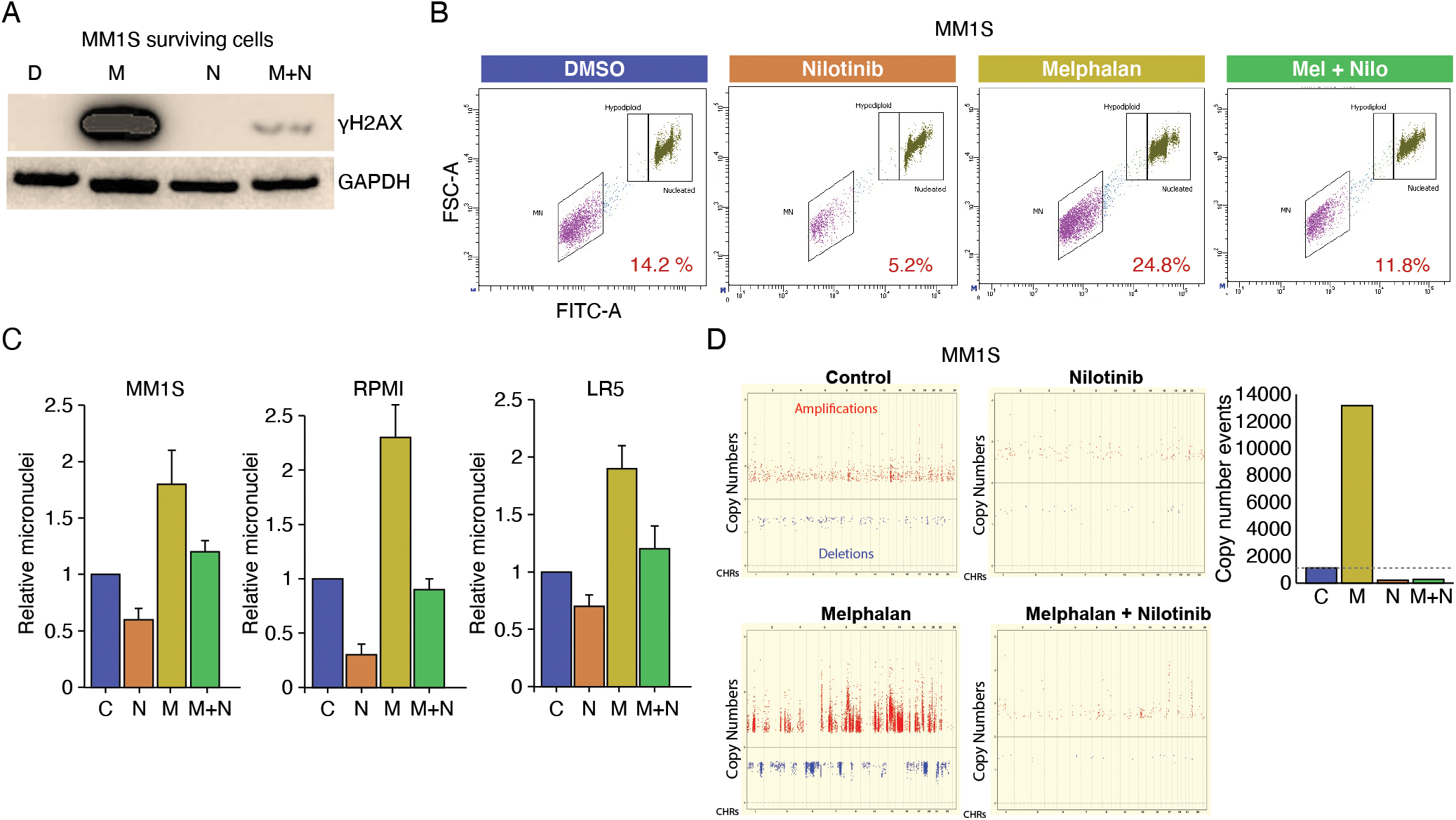
Nilotinib reduces genomic damage and instability caused by melphalan. **(A-B)** MM cells were treated with DMSO only (D), melphalan (M), nilotinib (N) or a combination of both drugs (M+N) for 48 hrs and evaluated for DNA breaks (by measuring γ-H2AX level) **(A)** and genomic instability (using micronucleus assay) **(B)**. A representative flow cytometry image of micronuclei in MM1S cells and **(C)** bar graphs showing percent micronuclei (with error bars representing SDs of three independent experiments) in three MM cell lines are shown. **(D)** MM1S cells treated with nilotinib (N; 2.5 μM), low dose melphalan (M; 1 μM) or combination were cultured for 3 weeks. DNA from the cultured and “day 0” (baseline control) cells was analyzed using SNP arrays (Affymetrix). Copy number events in cultured cells were identified using the genome of “day 0” cells as a baseline. A copy number event was defined as a change in ≥ 3 consecutive probes by 1 copy. Images showing amplifications (red dots) and deletions (blue dots) on all chromosomes and bar graph showing total copy number change events throughout genome.

### Inhibition of ABL1 reduces MM cell viability and increases cytotoxicity of chemotherapeutic agents in MM cells

Since ABL1 inhibition prevents a cancer cell from attempting to repair its DNA through HR, we next asked whether ABL1 inhibition reduces cell viability. We inhibited ABL1 in MM cell lines (MM.1S and RPMI8226) using shRNA and found significantly reduced viability of both MM cell lines (**Figure 5A**). Furthermore, ABL1 knockdown significantly increased the cytotoxicity of melphalan in both MM cell lines tested (**Figure 5B**). Consistently, nilotinib also reduced MM cell growth in a dose-dependent manner, with minimal or no impact on three different normal/non-cancerous cell types (HS5, human bone marrow stromal cells; HDF, human normal diploid fibroblasts; PBMC, peripheral blood mononuclear cells from two different donors). Nilotinib also sensitized three MM cell lines, including a melphalan-resistant cell line (LR5), to melphalan treatment with a combination index < 1, indicating a synergistic impact (**Figure 5C, D**).

**Figure 5:**
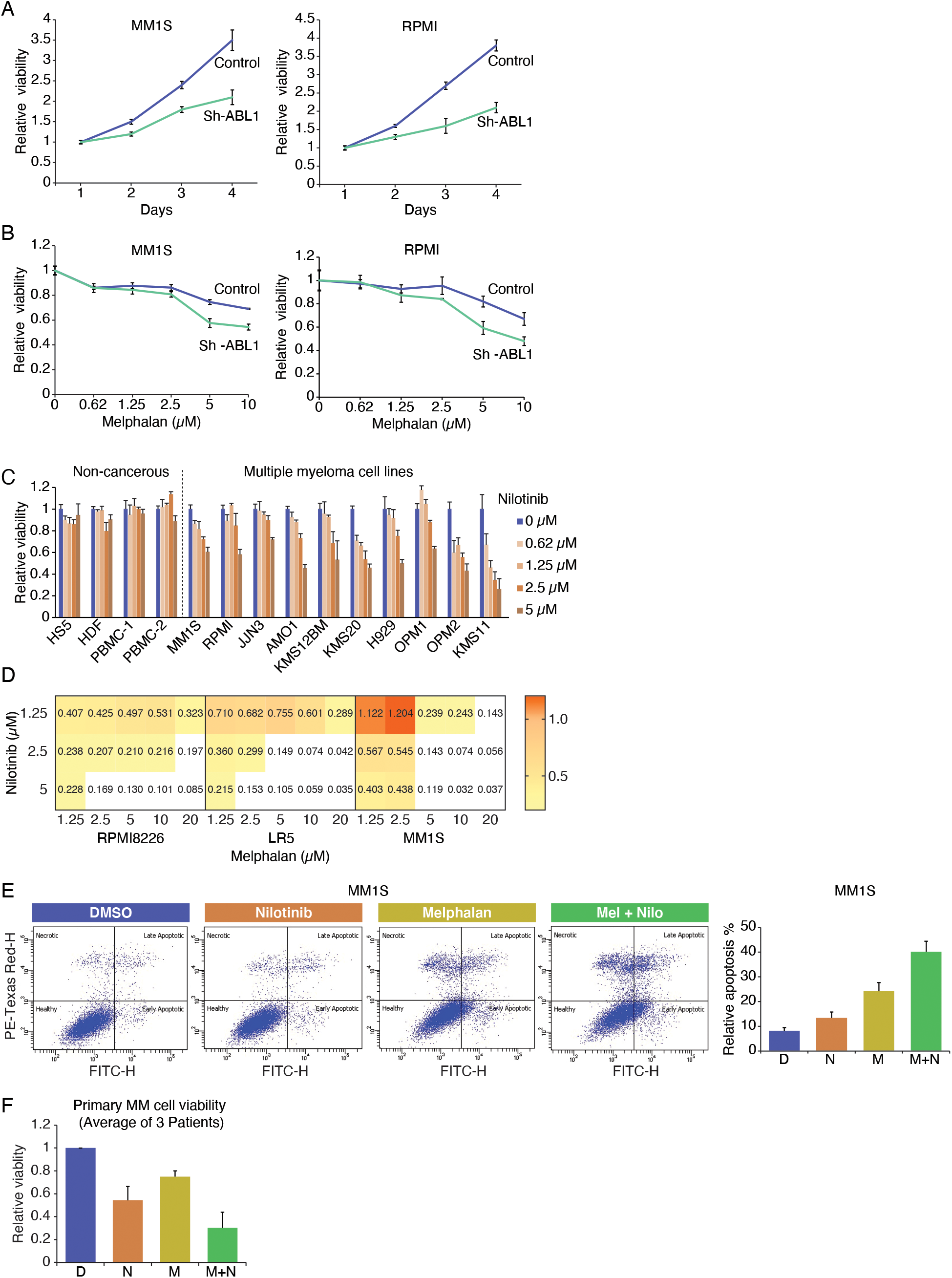
ABL1 inhibition reduces MM cell viability and increases cytotoxicity of melphalan in MM cells. **(A)** ABL1 in MM cells was inhibited using shRNAs (control, C; ABL1-targeting, sh-ABL1) and, following selection, cell viability was assessed for 4 days. **(B)** Control and ABL1-knockdown (sh-ABL1) MM cells were treated with different concentrations of melphalan for 48 hrs and cell viability measured. **(C)** A panel of MM cell lines were treated with different concentrations of nilotinib, and cell viability was measured. Bar graph showing impact of nilotinib on MM cell viability; error bars represent SDs of triplicate assays. **(D)** MM cell lines (MM1.S, RPMI8226 and LR5) were treated with different concentrations of nilotinib and melphalan, and cell viability measured after 48 hrs and combination index calculated using CalcuSyn software. (**E)** MM1S cells were treated with either DMSO (D), 2.5 μM nilotinib (N), 5 μM melphalan (M) or a combination (M+N), and evaluated for apoptosis using annexin/PI staining. Flow cytometry images of a representative experiment and bar graphs showing apoptosis in triplicate experiments are shown. **(F)** Bone marrow plasma cells from 3 relapsed MM patients were treated as indicated for 24 hrs and cell viability measured; error bars represent SDs of triplicate assays.

Flow cytometry–based annexin/PI staining demonstrated that treatment of MM.1S cells with nilotinib, melphalan and a combination of both drugs was associated with apoptosis in ~12%, 25% and 40% of cells, respectively, indicating that nilotinib increases melphalan-induced apoptosis in MM cells (**Figure 5E**). We also evaluated the impact of other DNA damage–related chemotherapy drugs (bendamustine and PARP inhibitor olaprib) in combination with nilotinib in MM cell lines and found nilotinib also significantly sensitized MM cells to these chemotherapeutic drugs (Supplementary Figures 1–2). Importantly, nilotinib also significantly sensitized bone marrow plasma cells from relapsed MM patients to melphalan treatment (**Figure 5F**).

### Inhibition of ABL1 significantly increases cytotoxicity of melphalan in a subcutaneous model of MM

We next evaluated the impact of nilotinib, melphalan and their combination in an *in vivo* subcutaneous tumor model of MM. Briefly, MM.1S cells were injected subcutaneously into SCID mice and, following the appearance of tumors, mice were treated with nilotinib (50 mg/kg; 5 days per week) and/or melphalan (2 mg/kg; twice per week) and tumor volumes measured regularly. After two weeks of treatment, tumor volumes in mice treated with vehicle control, nilotinib, melphalan and a combination of both drugs increased by 6-fold, 3-fold, 2.8-fold and 1.2-fold, respectively (P ≤ 0.02), relative to their initial sizes. The combination treatment resulted in the smallest tumor volumes in mice (**Figure 6A**), demonstrating the ability of nilotinib to increase the cytotoxicity of melphalan *in vivo*. At the end of two weeks, mice were euthanized, and the tumors removed and evaluated for HR activity in the lysates of the tumor cells. Consistent with *in vitro* data, melphalan treatment was associated with a significant increase in HR activity, whereas nilotinib significantly inhibited both the endogenous and melphalan-induced HR activities *in vivo* (**Figure 6B**).

**Figure 6:**
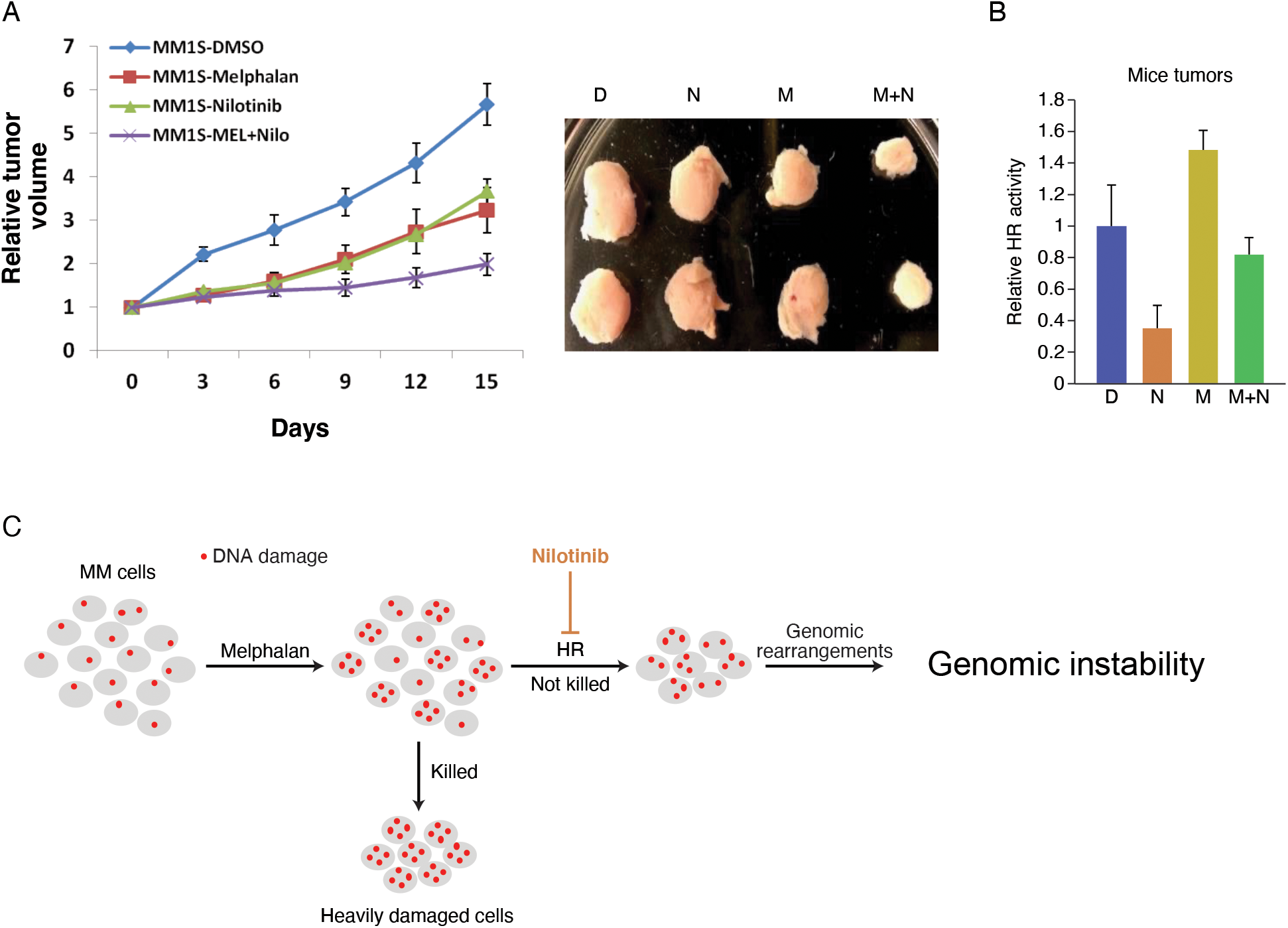
Nilotinib increases melphalan sensitivity in MM cells *in vivo*. **(A)** MM1S cells were injected subcutaneously into SCID mice and, following the appearance of tumors, mice were treated with either DMSO, nilotinib (50 mg/kg, 5 days per week), melphalan (2 mg/kg, twice per week) or in combination. Line plots showing tumor growth and images of representative tumors in different treatments are shown. **(B)** Tumors were lysed and evaluated for HR activity by *in vitro* strand exchange assay; error bars represent SDs of triplicate assays. **(C)** Model explaining the impact of nilotinib on melphalan-induced cytotoxicity and genomic instability. Melphalan treatment increases DNA damage, and the cells with very high levels of damage undergo apoptosis. However, cells with less damage survive. Increased HR activity in these cells not only helps in their survival but also contributes to aberrant repair and genomic instability. The addition of nilotinib inhibits HR, thereby increasing cell death and reducing genomic instability.

These data suggest that nilotinib increases the cytotoxicity of chemotherapeutic agents, such as melphalan, while reducing genomic instability in multiple myeloma cells **(Figure 6C)**.

## Discussion

Dysregulation of HR induces imprecise, unscheduled and unnecessary recombination events leading to genomic rearrangements and instability^21–27^. This genomic instability arises early during oncogenesis and contributes to oncogenic transformation and the progression of cancer^1–10^. For example, in esophageal cancer, elevated HR not only mediates genomic instability but also contributes to telomere maintenance and tumor growth *in vivo*^28^. Targeting mechanisms that promote genomic instability may be an effective strategy to delay progression, prevent the development of resistance to treatment, minimize the harmful genomic impact of chemotherapy and increase the cytotoxicity of chemotherapeutic agents. As such, drugs (such as nilotinib) that inhibit HR may delay progression in MM patients and other cancers. In fact, a clinical trial for CML shows that 99% of the patients treated with nilotinib did not progress to the accelerated phase.

One major problem in managing cancer patients is the development of resistance to treatment. The mechanisms underlying this problem could be intrinsic to cancer cells, acquired during treatment, or induced by treatment itself. These mechanisms may include altered cellular drug transport, inadequate cytotoxicity of the therapeutic agent, alterations in the drug targets and altered DNA repair pathways. Most chemotherapeutic agents induce DNA breaks, thus killing cells. However, surviving cells have more DNA breaks and higher HR activity, predisposing them to genomic instability and eventual genomic evolution towards drug resistance. Here, we found that cells surviving melphalan treatment had increased γ-H2AX expression, and this increase was blocked (to near background level) by the addition of nilotinib. Consistently, the genomic instability was also increased by melphalan and prevented by the addition of nilotinib. Importantly, these analyses were done on live cell fractions only, as apoptosis is associated with DNA breaks. Therefore, in the presence of dead cells, an accurate interpretation of γ-H2AX results is not possible.

Although melphalan-induced DNA breaks and genomic instability in MM cells were reduced by nilotinib, its cytotoxicity was increased. This may seem unexpected; however, myeloma cells have high endogenous DNA damage and overactive HR, which contributes to ongoing genomic rearrangements and instability^11,27^ as well as the growth and survival of cancer cells^28^. In this environment, melphalan treatment further increases DNA damage as well as HR activity, thus prompting heavily damaged cells to undergo apoptosis. However, in surviving cells, dysregulated HR not only helps them survive (by engaging the breaks) but also increases genomic rearrangements and instability.

In summary, our study is the first to demonstrate that a clinically approved ABL1 inhibitor (nilotinib) has the potential to inhibit growth and reduce genomic evolution in MM cells. When used in combination with a chemotherapeutic agent such as melphalan, nilotinib can reduce chemotherapy-induced genomic damage and instability, while increasing its cytotoxicity. Such treatments also have the potential to delay or prevent disease progression including development of drug resistance in myeloma.

## Acknowledgements

This work was supported by Department of Veterans Affairs Merit Review Award I01BX001584-01 (NCM), NIH grants P01-155258 and P50 CA100707 (NCM, MAS) and Paula and Rodger Riney grant (NCM).

## Author contributions

**SK** executed major experiments, analyzed and interpreted the data and prepared the manuscript; **ST, JZ, CL, LBP, SM, JS, CC, YT** carried out specific experiments and assisted in data analysis; **LB, MKS** analyzed SNP data and helped with statistical and bioinformatic analyses; **MAS** envisioned the study and participated in designing, data interpretation, and preparation of the manuscript; **NCM** envisioned the study and participated in its design and coordination, and also helped in drafting the manuscript. All authors have read and approved the final manuscript.

