## Supplemental Figures for "ABL1 Kinase plays an important role in spontaneous and chemotherapy-induced genomic instability in multiple myeloma"

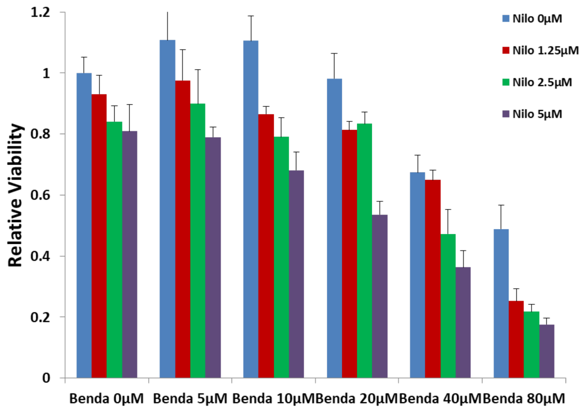


**Supplementary Figure 1. Nilotinib sensitizes MM cells to bendamustine.** MM cells were treated with different concentrations of nilotinib and bendamustine for 48 hrs, and the cell viability was measured.

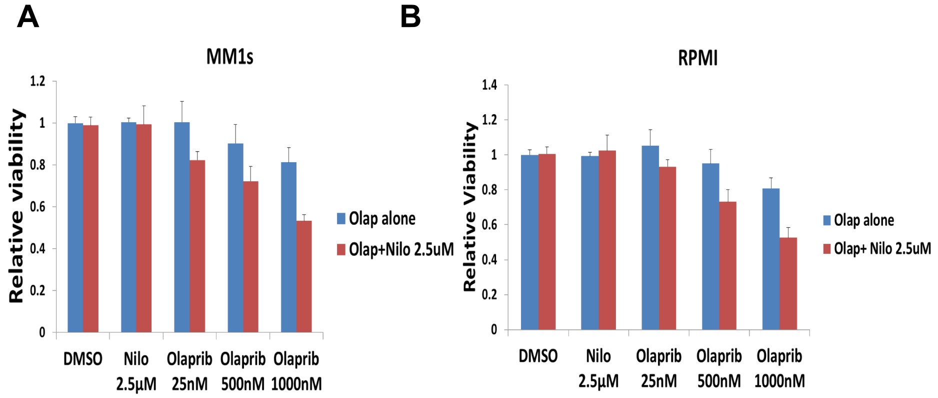


**Supplementary Figure 2. Nilotinib sensitizes MM cells to PARP inhibitor.** MM cell lines, MM1S (A) and RPMI8226 (B), were treated with PARP inhibitor at different concentrations, alone or in the presence of nilotinib (2.5 μM) for 48 hrs, and cell viability was measured.
